# A minimum Bayes Factor based threshold for activation likelihood estimation

**DOI:** 10.1101/2022.08.02.502470

**Authors:** Tommaso Costa, Donato Liloia, Franco Cauda, Peter Fox, Francesca Dalla Mutta, Sergio Duca, Jordi Manuello

## Abstract

Activation likelihood estimation (ALE) is among the most used algorithms to perform neuroimaging meta-analysis. Since its first implementation, several thresholding procedures had been proposed, all referred to the frequentist framework, returning a rejection criterion for the null hypothesis according to the critical p-value selected. However, this is not informative in terms of probabilities of the validity of the hypotheses. Here, we describe an innovative thresholding procedure based on the concept of minimum Bayes factor (mBF). The use of the Bayesian framework allows to consider different levels of probability, each of these being equally significant. In order to simplify the translation between the common ALE practice and the proposed approach, we analised six task-fMRI/VBM datasets and determined the mBF values equivalent to the currently recommended frequentist thresholds based on Family Wise Error (FWE). Sensitivity and robustness toward spurious findings were also analyzed. Results showed that the cutoff log_10_(mBF)=5 is equivalent to the FWE threshold, often referred as voxel-level threshold, while the cutoff log_10_(mBF)=2 is equivalent to the cluster-level FWE (c-FWE) threshold. However, only in the latter case voxels spatially far from the blobs of effect in the c-FWE ALE map survived. Therefore, when using the Bayesian thresholding the cutoff log_10_(mBF)=5 should be preferred. However, being in the Bayesian framework, lower values are all equally significant, while suggesting weaker level of force for that hypothesis. Hence, results obtained through less conservative thresholds can be legitimately discussed without losing statistical rigor. The proposed technique adds therefore a powerful tool to the human-brain-mapping field.

## Introduction

Since the seminal work of Ogawa, Lee, Kay, and Tank (1990), research based on magnetic resonance imaging (MRI) has produced a large amount of data associated to cognitive domains and sensory processes, in both healthy and diseased populations. However, the high complexity of the topic investigated, together with peculiarities of the used techniques, expose results to a potential high variability (Bowring, Maumet,& Nichols, 2019; Smith et al., 2005). This issue can be overcome by means of coordinate-based meta-analysis (CBMA), a technique allowing to pool results from different experiments that investigated similar questions (Wager, Lindquist, & Kaplan, 2007; Wager, Lindquist, Nichols, Kober, & Van Snellenberg, 2009). Notably, this allows to produce additional statistics based on a much greater sample size of those analyzed in the original experiments, which sometimes lead to scarcely reliable individual results (Manuello, et al., 2022).

Among the available techniques to perform CBMA, activation likelihood estimation (ALE) aims to estimate spatial convergence of activation probabilities between the results of previously published experiments. It is based on the null-hypothesis that the foci (i.e., the peaks of effect reported in each single experiment) are uniformly spread across the brain, while looking for regions in which the convergence across multiple experiments is higher than if it came from an independent distribution (Eickhoff, Bzdok, Laird, Kurth, & Fox, 2012). One of the crucial elements of ALE is the thresholding procedure, and for this reason it evolved over time to improve statistical reliability of the results obtained by means of this meta-analytic approach. In the first implementation, the voxel-wise significance was assessed based on a fixed-effect analysis without correction for multiple comparisons. Since the lack of this correction was likely to introduce false positives, successive modifications introduced the false discovery rate (FDR) approach to account for this (Laird et al., 2005). Later on, Eickhoff et al. (2009) empirically estimated the spatial variability of data for the Gaussian filtering and optimized the implementation of the permutation test. Finally, Eickhoff et al. (2012) integrated the method with two additional thresholding procedures based on the correction for family-wise error (FWE) rate and the cluster-level significance (c-FWE). All these inferential procedures are theoretically based on the conjoint use of Fisher’s p-value approach and Neyman-Pearson’s Test approach, which are often combined in the scientific practice.

Here, we aimed to introduce an innovative thresholding option, based on Bayesian inference rather than on the frequentist one. Specifically, this leverages on the concept of minimum Bayes Factor (mBF), which is the smallest Bayes Factor obtained for a p-value in a given distribution with respect to the alternative hypothesis. The mBF has several advantages over the available canonical thresholding procedures. On the statistical side, the canonical ALE-related thresholds, which are based on frequentist inference, solely provide a rejection criterion for the null hypothesis according to the critical p-value selected. However, this is not informative in terms of probabilities of the validity of hypotheses (Costa et al., 2021). On the contrary, the Bayesian solution allows to consider different levels of probability, each of these being equally significant. Moreover, the BF, that is a unique function of the data, allows a straightforward interpretation, as it expresses how probable it is to find a real effect in a given voxel. Secondarily, on the computational side, it does not require any random or permutation test. This is particularly relevant when analyzing large datasets of experiments, allowing to significantly reduce the required processing time (i.e., a few seconds versus tens of minutes, up to hours in worst cases).

In the present work, we first provided a formal description of the BF and the mBF. Then, aiming to facilitate the comparison between the common ALE practice and the proposed new procedure, we analyzed a range of meta-analytic datasets to identify the mBF equivalent to canonical thresholds. Based on these values, we compared the sensitivity of the mBF with that of other available solutions. Finally, we tested the robustness of the Bayesian approach towards false positives detection. The introduction of this innovative thresholding procedure could therefore further consolidate one of the most used methodologies in CBMA research, allowing a deeper interpretation of the results, while simplifying at the same time its computational implementation.

## Materials and methods

### Anatomical likelihood estimation and creation of the modeled activation maps

An in-depth description of the ALE methodology (Eickhoff et al., 2012; Eickhoff et al., 2009; Turkeltaub et al., 2012) is out of scope for the present work. Here, we briefly recapitulate its core concepts following the nomenclature used in Samartsidis, Montagna, Nichols, and Johnson (2017). In ALE environment, a Gaussian distribution is centered on each voxel, representing the probability to find a true effect in it. In other words, this means computing the likelihood of the voxel to be the exact location of a focus reported in one (or more) of the original experiments included in the dataset. Given the experiment *i*, the map based on one of its focus *x_ik_* is expressed as:

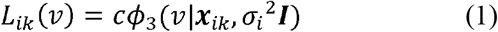

where *ϕ*_3_(*x*; *μ*,∑) is a three-dimensional Gaussian distribution with mean and covariance *μ*, ∑ evaluated in 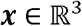. *c* is a normalization constant ensuring that the sum of *ϕ*_3_ over voxels equals to 1, and *I* is the identity matrix. Therefore, *L_ik_*(*ν*) represents the probability that the voxel *ν* is the true location of the focus ***x**_ik_*. The only free parameter in formula 1 is *σ_i_*. Its value was empirically determined in Eickhoff et al. (2009), considering the number of subjects *n_i_* in each experiment.

After this first step, the *L_ik_* maps obtained for each single focus reported in a given experiment are combined in *L_i_* also named modelled activation (MA) map. Each MA represents the estimate of the full effect map obtained in the original experiment and described in a reduced form through the associated list of foci. Finally, the ALE statistic is computed as follows:

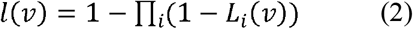

This formula represents the probability for the voxel *ν* to be the location of a true effect coherently measured across the experiments included in the dataset. This algorithm is implemented in GingerALE software (https://www.brainmap.org/ale/) (Eickhoff et al., 2012; Eickhoff et al., 2009; Turkeltaub et al., 2012). In addition to the unthresholded and thresholded ALE maps, the output also includes a Z map. The conversion from p-values to Z points is based on the numerical method proposed by Cody (1969).

### Bayes Factor and the minimum Bayes Factor

The computation of the mBF is based on the Bayes’ theorem. According to it, the probability associated with two concurrent hypotheses, namely *H*_0_ and *H*_1_, can be formalized as follows:

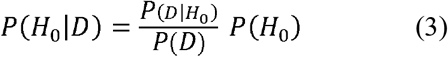

and, correspondingly,

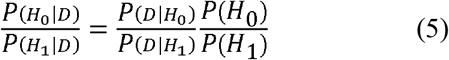

where *D* is the measured effect in a voxel.

Their quotient represents the Bayes’ theorem in terms of relative belief:

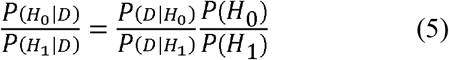

Based on the previous formulas, the *BF*_01_ can therefore be expressed as:

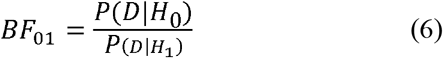

The value of *BF*_01_ gives the degree of evidence for the two concurrent hypotheses: when *BF*_01_ > 1 the evidence favors *H*_0_; on the contrary, when *BF*_01_ < 1 the evidence favors *H*_1_.

In the usual form, using the Bayes Factor to measure the evidence for a composite hypothesis requires averaging all possible distinct alternative values. However, in some cases infinite alternative values exist, as for instance for the hypothesis “the mean is different from zero”. Therefore, determining a point value among all the possible ones of the composite hypothesis can be extremely complex and sometime impractical. To circumvent this issue, it is possible to select the alternative value with the largest effect against the null hypothesis, considering this as the summary of evidence across all the possible distinct values contained in the composite alternative hypothesis. This is called the Minimum Bayes’ Factor (mBF), which therefore reflects the largest amount of evidence against the null hypothesis. In other words, the mBF uses the best-supported alternative hypothesis *H*_1_ This represents the worst-case scenario for *H*_0_ among all possible distinct Bayes’ factors values, because no alternative value is supported by a larger amount of evidence against *H_0_* than the one identified through the mBF.

Let’s now consider as the hypothesis the probability of a generic observed effect *x* modelled through a Gaussian distribution with mean *μ* and variance *σ*^2^:

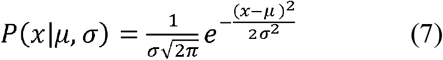

the BF for the null hypothesis versus the supported hypothesis (*μ* = *x*) is the mBF:

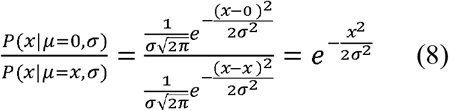

If we now take the Z map obtained from the unthresholded ALE map, the mBF can be written as:

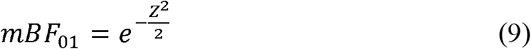

by applying a simple exponentiation. This represents the evidence of the null hypothesis *H*_0_ against the alternative hypothesis *H*_1_. To obtain the evidence of *H*_1_ the reciprocal of the *mBF*_01_ can be computed as:

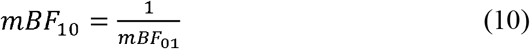

The mBF ranges from 0 to ∞, and according to Kass and Raftery (1995) values can be interpreted as shown in Table 1. The MATLAB^®^ code which implements the Z to mBF transformation described in formula 10 is freely available on figshare (https://figshare.com/articles/software/minimum_bayes_factor_script/17023931).

**Table 1.** The evidence categories for the Bayes Factor (BF). Adapted from Kass and Raftery (1995).

| BF value | Force of evidence |
| --- | --- |
| 1 - 3 | Very weak |
| > 3 - 20 | Positive |
| > 20 - 150 | Strong |
| > 150 | Very strong |

### Identification of the equivalent mBF thresholding

Since its inception, the ALE has been based on frequentist statistics, and researchers applying it are therefore used to reason in frequentist terms. Therefore, in order to facilitate the mental translation between the p-value based interpretation and Bayesian one, we first aimed to find mBF values equivalent to canonical thresholds applied to ALE. To do so, the ALE algorithm was applied to six different datasets, three of them referring to task-related functional MRI (fMRI) data, the remaining three including only voxel-based morphometry (VBM) experiments. In order to maximize reliability and reproducibility, five of these datasets were selected from published CBMAs based on ALE method. Briefly, the first task-fMRI pool consisted of 109 experiments on time-perception, originally collected and analyzed in Nani et al. (2019). A second functional pool included 73 experiments based on finger-tapping task, from Laird et al. (2008). The third and last one gathered 19 experiments on face discrimination, from Eickhoff et al. (2012). On the structural side, the largest pool included 57 experiments on Alzheimer’s disease, previously analyzed in Manuello et al. (2018). A further pool focused on chronic schizophrenia, with 49 experiments collected in Liloia et al. (2021). Finally, the last pool was built querying the VBM section of the BrainMap database (Vanasse et al., 2018) to retrieve 20 available experiments on mild cognitive impairment (MCI) (see also Table S1 in the Supplement). It should be noted, that the selected datasets had varying sizes. This was decided in order to verify the possible confounding effect of the number of experiments analysed. In particular, the face discrimination and MCI datasets were selected to verify the behavior of mBF threshold when the sample size approaches the lower bound recommended for valid ALE analyses (Eickhoff et al., 2016).

The identification of the equivalent mBF threshold was then based on a four steps procedure, separately applied to each dataset: i) Data were analyzed through GingerALE (v.3.0.2), applying different canonical thresholds with the default settings suggested by the tool. To be more precise, for the fixed-effect uncorrected thresholding the p-value was set to 0.05. For the cluster-level thresholding the uncorrected p-value was set to 0.001, the corrected p-value to 0.05, tested with 1000 permutations. For the voxel-level thresholding the corrected p-value was set to 0.05, tested with 1000 permutations. ii) The thresholded ALE maps obtained in the previous step were converted into binary masks. iii) The Z map computed by GingerALE was converted into a BF map, as shown in formulas 9-10. iv) The BF map was individually masked with each of the previously binarized thresholded ALE maps. At this point, the lowest voxel value in the obtained output was taken as the mBF threshold. In other words, this is the cutoff that can be set to apply the Bayesian equivalent of the canonical threshold. In terms of interpretation, the BF values in the map express how much more probable it is to find a true convergence of effect in a given voxel as compared to finding no convergence. In order to prove its reliability, the mBF equivalent to a given canonical threshold must be consistent across the different datasets analysed.

### Correlation between BF and canonical ALE maps

For the purpose of further investigating the behavior of the Bayesian approach, we aimed to identify the mBF value that maximizes the correlation with the ALE maps canonically thresholded. In details, ALE maps were computed, thresholded and binarized as described above in steps i and ii. The BF map was obtained as in step iii, and then expressed in logarithmic scale. This log_10_(BF) map was iteratively binarized applying increasing threshold values, and Pearson correlation was run with the binarized ALE map at each step. Notably, the mBF previously obtained matches the mBF returning the maximum correlation only if the mBF-thresholded map maximizes the overlap with the canonical ALE map, while minimizing non-overlapping voxels.

### Sensitivity of the equivalent mBF thresholding

As previously described, the mBF threshold was identified by means of masking procedure base on the canonical ALE maps. However, this does not exclude in a non-masked map the presence of further voxels with the same mBF value but non-overlapping with the canonical ALE map of reference. Therefore, in order to be a suitable alternative to the frequentist thresholds, the mBF must produce maps covering the highest possible fraction of the canonical ALE map, while minimizing non-overlapping voxels. Moreover, the potentially remained non-overlapping voxels should be localized close to the clusters of overlapping voxels, rather than randomly spread throughout the brain. These can be qualitatively assessed by a visual comparison of the mBF and canonical ALE maps. To obtain a quantitative measurement, the log_10_(mBF) map was iteratively binarized at increasing threshold values as described above, and at each step the number of voxels appearing only in the mBF map, only in the ALE map, or in both of them was counted. Finally, the log_10_(mBF) threshold allowing to remove any voxel appearing in the Bayesian map but not in the canonical ALE map was computed.

### Robustness towards spurious convergences

Since the ALE method aims to identify spatial convergence among multiple experiments, the frequentist thresholds were designed to suppress possible spurious convergences in the results. As the same behavior is required from the Bayesian approach, we tested it on a dataset including 20 experiments that investigated unrelated cognitive domains (see Table S2 for a description of this dataset), and on two simulated datasets generated with the Fail-safe R script (Acar et al. (2018); https://github.com/NeuroStat/GenerateNull) and built with 21 experiments each with foci randomly placed across the brain. In this condition, possible voxels surviving the threshold must be considered as spurious evidence, therefore the mBF approach is expected to produce empty maps.

## Results

### Identification of the equivalent mBF

With the aim of identifying an equivalent mBF value for the different canonical thresholds, six datasets were processed with ALE algorithm and thresholded with both Bayesian and canonical frequentist approaches. Results showed a consistent behavior of the mBF (Table 2). In details, the p05 threshold was shown to be equivalent to a mBF of ≈4. As shown in Table 1, this is a rather weak evidence, in line with the already known low reliability of its equivalent frequentist solution. Since the use of p05 thresholding is to date explicitly discouraged in the common ALE practice (Eickhoff et al., 2016), no further analyses were carried on for its Bayesian counterpart. The c-FWE threshold was shown to be equivalent to a mBF of ≈119. Finally, the FWE threshold was shown to be equivalent to a mBF of ≈10^5^. Notably, the equivalent mBF value for c-FWE and FWE can be interpreted as ‘strong’ and ‘very strong’ evidence respectively. No differences were found between task-fMRI and structural datasets, and no effect of sample size was highlighted. The iterative correlation between the maps thresholded with either the Bayesian approach or the canonical one confirmed the mBF values previously identified. In most cases peaks exceeded r = 0.9, considering either FWE or c-FWE (Fig. 1). Interestingly, in the two datasets with sample size around the recommended minimum (i.e., MCI and face discrimination), the correlation peaks with FWE were considerably higher than those obtained with c-FWE.

**Table 2.** mBF values equivalent to canonical frequentist thresholds. In the last two columns, the maximum Pearson correlation value obtained between the Bayesian and canonical ALE maps is shown (this information is also graphically represented in **Fig.1**). Values in brackets refer to the number of experiments in the dataset.

| MRI Modality | Dataset | equivalent minimum_BF |  |  | max correlation |  |
| --- | --- | --- | --- | --- | --- | --- |
|  |  | p05 | FWE | c-FWE | FWE | c-FWE |
| Task fMRI | Time perception (109) | 3.8711 | 1.97E+05 | 1.19E+02 | 0.95 | 0.95 |
|  | Finger tapping (73) | 3.8816 | 2.12E+05 | 1.19E+02 | 0.96 | 0.98 |
|  | Face discrimination (19) | 3.8891 | 1.13E+05 | 1.19E+02 | 0.99 | 0.90 |
| VBM | Alzheimer (57) | 3.8724 | 2.35E+05 | 1.19E+02 | 0.94 | 0.92 |
|  | Schizophrenia (49) | 3.8754 | 2.44E+05 | 1.19E+02 | 0.86 | 0.83 |
|  | MCI (20) | 3.872 | 9.96E+04 | 1.19E+02 | 0.98 | 0.85 |

**Fig. 1:**
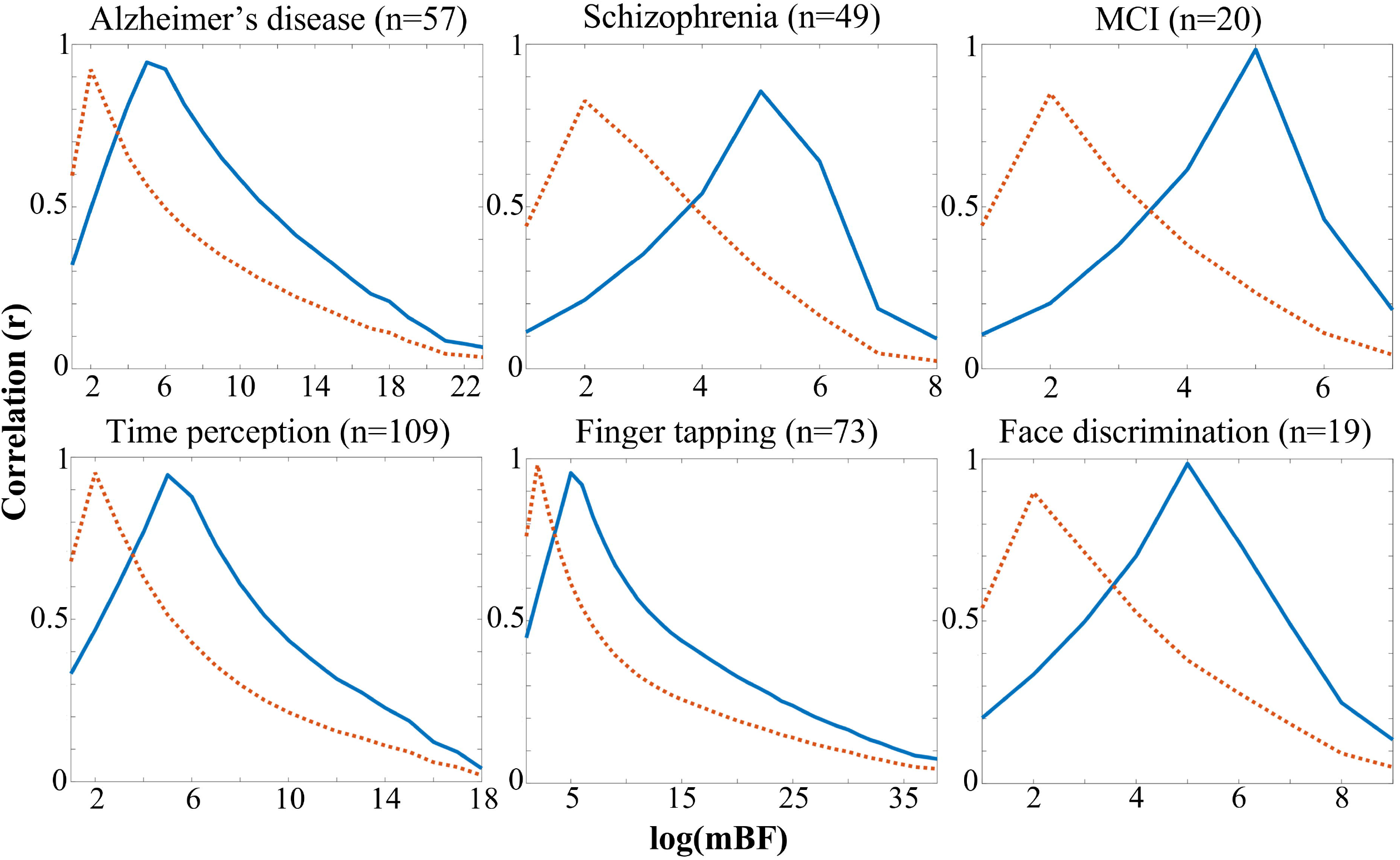
The iterative Pearson correlation between the maps thresholded with either the Bayesian approach or the canonical one. The blue line refers to the correlation with the FWE thresholding, while the orange dashed line refers to the correlation with the c-FWE thresholding. The x axis represents the log_10_(mBF) value used to threshold the Bayesian map. Values in brackets refer to the number of experiments in the dataset. MCI = mild cognitive impairment.

### Sensitivity and robustness

The qualitative comparison between the canonical ALE map and the equivalent mBF-thresholded map showed, in line with the observed correlation peaks, a substantial overlap between the two. However, the spatial distribution of the non-overlapping mBF voxels differed between FWE and c-FWE. For what concerns the former, the non-overlapping voxels surviving the threshold of log_10_(mBF)=5 where localized around the regions of overlap (Fig. S1). On the contrary, voxels surviving the threshold of log_10_(mBF)=2 in the latter case formed blobs far from the regions of overlap (Fig. S2). Notably, in order to suppress voxels present in the mBF map but not in the c-FWE ALE map, the log_10_(mBF) threshold had to be increased of orders of magnitude, being at least 4 for finger tapping and 8 for time perception (Table 3 and Fig. 2). However, this also removed up to the 94.5% of voxels in overlap, in the worst case. This loss of convergence was much reduced in the case of FWE ALE maps, never exceeding the 59%. Finally, it is relevant to note that in all cases the equivalent mBF threshold marks the end of the steady state of the number of overlapping voxels between the mBF map and the canonical ALE map. This same landmark also coincides with the correlation peak between the two conditions.

**Table 3.** log_10_(mBF) values necessary to suppress voxels present in the Bayesian map but not in the canonical ALE map, together with the percentage of overlapping voxel lost while increasing the Bayesian threshold from the equivalent mBF previously identified to the more stringent suppressing threshold. Values in brackets refer to the number of experiments in the dataset.

| MRI Modality | Dataset | FWE |  | c-FWE |  |
| --- | --- | --- | --- | --- | --- |
|  |  | non overlap suppression th | % of overlap lost | non overlap suppression th | % of overlap lost |
| Task fMRI | time_perception (109) | 6 | 22.8 | 8 | 91.0 |
|  | finger_tapping (73) | 6 | 15.2 | 4 | 49.0 |
|  | face_discrimination (19) | 6 | 44.7 | 5 | 85.6 |
| VBM | Alzheimer (57) | 6 | 14.7 | 6 | 75.3 |
|  | Schizophrenia (49) | 6 | 59.0 | 5 | 91.0 |
|  | MCI (20) | 5 | 3.3 | 5 | 94.5 |

**Fig. 2:**
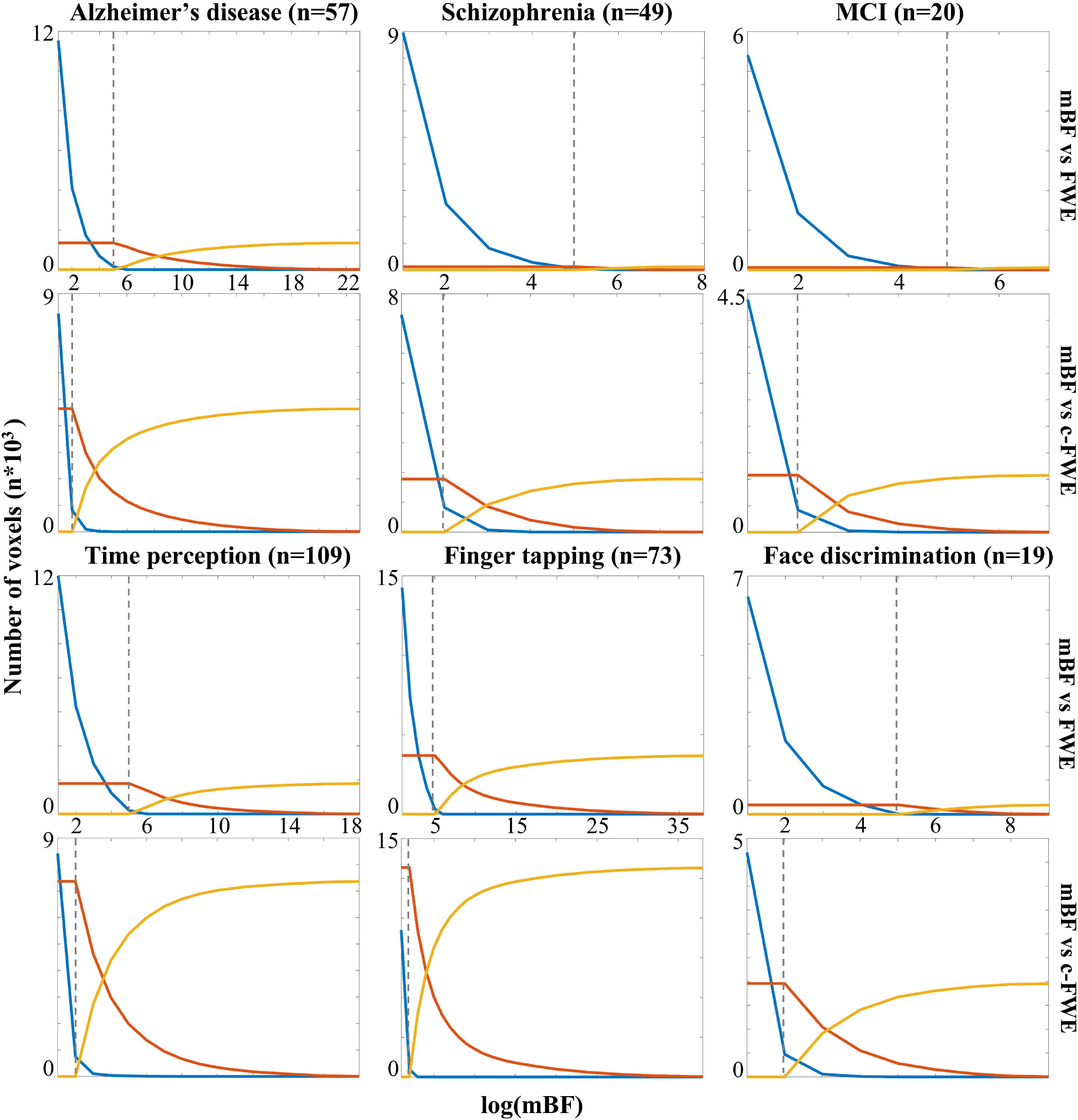
Number of non-zero voxels appearing only in the mBF-thresholded map (blue line), only in the canonically-thresholded map (yellow line), or in overlap between the two maps (red line). The x axis represents the log_10_(mBF) value used to threshold the Bayesian map. The vertical dashed line marks the log_10_(mBF) threshold equivalent to canonical thresholds as previously identified. Values in brackets refer to the number of experiments in the dataset. MCI = mild cognitive impairment.

The analyses of the random tasks dataset and of the two random foci datasets confirmed that a log_10_(mBF)=2 is not conservative enough to suppress spurious convergences. On the contrary, no voxels in the maps reached a value of log_10_(mBF)=5 (See Fig. S3).

## Discussion

In the present work, we described and applied an innovative thresholding approach for ALE meta-analyses based on the mBF. The adoption of the Bayesian framework is a relevant development for the CBMAs field, which had included so far only the frequentist paradigm. In fact, although these two statistical families often tend to produce convergent findings (Jaynes & Kempthorne, 1976), they differ in the logic used to interrogate data and interpret results (Cauda et al., 2020; Costa et al., 2021; Poldrack, 2006). For this reason, while suggesting a new framework we first aimed to identify its quantitative equivalent in the frequentist counterpart. This consistently emerged across the six heterogeneous meta-analytic datasets analysed, showing the mBF≈4 cutoff being tantamount to the p05 canonical threshold, the mBF≈119 cutoff being tantamount to the c-FWE canonical threshold and the mBF≈10^5^ cutoff being tantamount to the FWE canonical threshold.

Although the interpretation of BF values proposed by Kass and Raftery (1995) was conceived for a general scope and is therefore not specific for neuroimaging results, it is possible to note that the force of the evidence retrievable through the equivalent of the p05 thresholding is “very weak”. On the contrary, results obtained with mBF≈119 (i.e., c-FWE) are scored as “strong”, and those obtained with mBF≈10^5^ (i.e., FWE) are “very strong”. Notably, Kass and Raftery’s cutoff for “very strong” evidence is BF>150, while values in our results are at least three orders of magnitude greater. This information, that was not obtainable through the frequentist approach, reveals that the ALE technique can produce extremely robust results, in the Bayesian meaning. Moreover, it seems to suggest the canonical FWE thresholding being more conservative than the c-FWE option. It is relevant to note that when ALE is applied to unrelated or random data, results did not reach the same strength of evidence observed for real datasets.

Since one of the advantages of ALE is to be adequate for both functional and structural MRI data, describing both the healthy or the pathological brain (Eickhoff et al., 2012) the consistent behavior of the mBF thresholding observed across all the tested datasets was strongly necessary. Similarly, the mBF approach showed to be reliable for the whole range of sample sizes being valid for ALE analysis (i.e., at least 17 experiments per dataset as recommended by Eickhoff et al. (2016)). However, contrary to the frequentist framework, the Bayesian approach doesn’t require a minimum number of evidences to produce unbiased results (Costa et al., 2021). For this reason, provided that every meta-analysis aims to include the largest possible fraction of the published literature on a given topic, the mBF thresholding now allows to perform CBMA in those specific domains which had been so far hampered by an insufficient number of retrievable experiments.

As a further advantage, the final ALE maps obtained through the mBF approach are easily comparable among them, as the values directly express the strength of the evidence. For example, the observed BF value of 200 in a voxel activated or altered by the ‘condition A’ would be exactly twice as strong of the BF value of 100 observed in a voxel activated or altered by the ‘condition B’. Extending the same linear relationship observed in the Bayesian case to the frequentist paradigm would be technically wrong. In fact, although sequentially developed over time, the p05, the FWE, and the c-FWE are based on different statistics (Eickhoff et al., 2016) and results obtained with them should not be interpreted as being lying on a continuum, despite the discussed trend of equivalent mBF values.

Similarly, the logic behind lowering the mBF threshold is not equivalent to the meaning of modifying the p-value. In fact, evidence obtained in the frequentist framework at p<0.01 is 100 times less accurate than evidence obtained at p<0.0001. On the contrary, evidence showing in the Bayesian framework a value of mBF=1 is 100 times weaker than evidence showing a value of mBF=100, but the two are equally accurate. The same difference should be considered when testing a-priori hypotheses. Let’s suppose, as an example, to expect to find the involvement of brain region A in a given cognitive domain, but then to not observe that in the final ALE map thresholded at FWE p<0.001. Increasing the p-value to 0.01 or even to 0.05 until the expected brain region appears in the map is a questionable procedure, that should be clearly highlighted by researchers. On the contrary, if that region doesn’t survive mBF=10^5^ it is licit to lower the threshold to whichever value, then discussing the observed strength for the hypothesis of its involvement in that cognitive domain. In light of this, there is no absolute cutoff to be recommended for the use of the mBF based ALE thresholding. However, the following standards should be followed to report and describe results obtained through mBF thresholding in a clear and transparent way: i) No binarization should be applied to the thresholded maps; ii) No upper-bound cutoff should be used to limit the range of values for visualization purpose (logarithmic scale can be used instead, if needed); iii) The maps should be always accompanied with a color scale clearly showing the range of values. If it is true that the mBF threshold can be adjusted, it is extremely relevant to know how the mBF values are distributed across the map. The implications of the recommended standards can be appreciated in figure 3. Applying the binarization, the small cluster in the left insula seems to be equally informative as the other two cluster, and it is impossible to localize any peak in the map. Similarly, the peaks remain hidden if setting an upper-bound cutoff. Conversely, the original map, arbitrarily thresholded at mBF=10^5^, allows to identify the peak of effect, although the rest of the maps appears flattened around the threshold. On the contrary, resorting to the logarithmic scale allows to appreciate the full distribution of values.

**Fig. 3:**
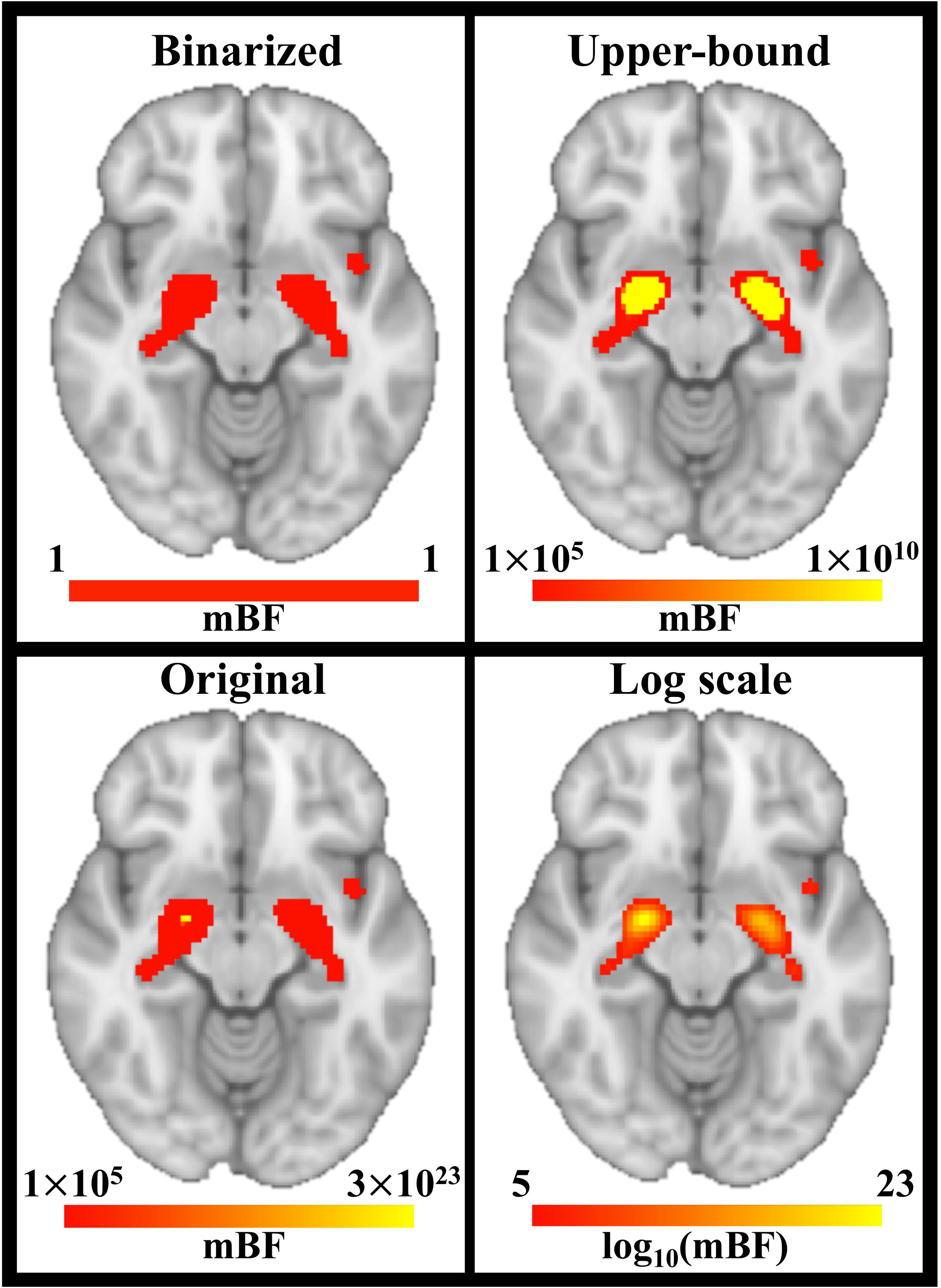
A visual representation of the recommended standards to report results in the Bayesian framework. In each condition the map was thresholded at mBF=10^5^. In the top left quadrant, the mBF=10^5^ threshold was also used to binarize the map. In the top right quadrant, an arbitrary upperbound cutoff at mBF=10^10^ was applied. The bottom left quadrant shows the original map. In the bottom right quadrant, the logarithmic scale was used instead. Results refers to the analysed dataset on Alzheimer’s disease. The axial slice is shown in radiological standard (left is right).

As previously mentioned, and following Kass and Raftery (1995), any evidence showing BF>150 should be considered as “very strong”. At the same time, our results showed voxels with value greater than log_10_(mBF)=35, being hardly comparable with Kass and Raftery’s range of 1-150. Therefore, as the standard practice in the ALE field has been so far based on frequentist thresholds, it sounds reasonable to highlight that the use of a mBF threshold of 10^5^ warranties results strongly comparable with those obtained through the FWE thresholding, generally considered a reliable standard (Eickhoff et al., 2016). On the contrary, a mBF threshold of 10^2^ shouldn’t be intended as tantamount to the c-FWE thresholding. This is likely ascribable to the lack of a topological criteria in the Bayesian approach, which is instead applied in the c-FWE algorithm after the initial uncorrected p-value estimate (Eickhoff et al., 2012). Finally, if compared with the computation of FWE-based thresholds in the context of ALE, mBF has the advantage of not being dependent on Monte-Carlo method, which is computationally intense and therefore potentially resulting in highly time-consuming analyses. While proposing an alternative procedure for statistical thresholding, the mBF approach builds on the ALE technique, and therefore inherits its limitations, as well as those of CBMAs in general. The main one pertains to the creation of the dataset, which should be as extensive as possible. Nevertheless, the Bayesian approach is less prone to the effects of potentially biased sampling, as it explicitly considers the specific set of data, without referring to any hypothetical whole population. As suggested by our results, the proposed approach should be equally efficient irrespective of the content of the domain analysed. While 6 datasets could be deemed as insufficient amount of data to provide reliable evidence in terms of generalizability, it should be noted that their selection was carefully designed to maximize their representativeness. We therefore included data describing both cognitive domains and pathologies, through functional or structural MRI data respectively. Moreover, the cognitive domains and related tasks were selected to be highly heterogeneous, and both psychiatric and neurodegenerative disorders were covered. As an additional point, the datasets were gathered between 2008 and 2021, therefore potentially capturing the effects of technical changes happening during the years. Finally, we also manipulated the number of experiments included in each dataset, which is commonly considered the factor to have more power to affect and bias results obtained through ALE. It is of course not possible to exclude a discrepancy of the equivalent mBF for canonical thresholds in specific datasets, but since there are no technical or theoretical reasons to expect markedly divergent behavior to happen systematically, there was no way to set an ideal amount of datasets to be analysed. Although the shared code to implement the transformation from Z maps to BF maps was written in MATLAB^®^, the same procedure can be adapted to any other language and programming environment. Finally, while we referred to the GingerALE software, any Z map produced as output of other implementations of the ALE algorithm is equally suitable for the mBF approach here described.

## Conclusions

CBMA is a widely used and meaningful technique in human neuroimaging. The adoption of a Bayesian framework for statistical thresholding, and specifically of the computation of mBF, can strengthen the ALE approach to CBMAs, expanding its field of application to small datasets, and further confirming the robustness of results obtained through this technique.

## Data availability statement

The data that support the findings of this study are available from the corresponding author upon reasonable request.

