## Supplementary material for "A minimum Bayes Factor based threshold for activation likelihood estimation"

Contents

Tables and Figure:

*Table S1.* Experiments included in the meta-analysis: Mild cognitive impairment dataset……p. 2

*Table S2.* Experiments included in the meta-analysis: Unrelated cognitive domains dataset….p. 3

*Figure S1.* Comparison between FWE thresholding and mBF thresholding …………………..p. 4

*Figure S2.* Comparison between c-FWE thresholding and mBF thresholding ……………….. p. 5

*Figure S3.* Results for unrelated cognitive domains dataset and simulated datasets …………..p. 6

*Table S1.* Experiments included in the meta-analysis: Mild cognitive impairment dataset.

| ***First author*** | ***Year*** | ***BrainMap ID*** | ***Journal*** | ***Subj*** | ***Foci*** |
| --- | --- | --- | --- | --- | --- |
| Chetelat G | 2002 | 8050043 | NeuroReport | 22 | 10 |
| Pennanen C | 2005 | 8050105 | Journal of Neurology, Neurosurgery, and Psychiatry | 32 | 10 |
| Bell-McGinty S | 2005 | 8060155 | Archives of Neurology | 4 | 7 |
| Bozzali | 2006 | 10010013 | Neurology | 20 | 14 |
| Hamalainen A | 2007 | 9010002 | Neurobiology of Aging | 14 | 6 |
| Karas G | 2008 | 9050041 | American Journal of Neuroradiology | 11 | 4 |
| Shiino A | 2006 | 9050064 | NeuroImage | 20 | 10 |
| Gold B T | 2010 | 11040071 | Human Brain Mapping | 12 | 4 |
| Barbeau E | 2008 | 11040158 | Neuropsychologia | 12 | 11 |
| Schmidt-Wilcke T | 2009 | 11040205 | NeuroImage | 18 | 4 |
| Rami L | 2009 | 11040232 | International Journal of Geriatric Psychiatry | 14 | 5 |
| Agosta F | 2011 | 13100085 | Radiology | 15 | 1 |
| Miettinen P S | 2011 | 13100144 | European Journal of Neuroscience | 18 | 5 |
| Guedj E | 2009 | 14080019 | European Journal of Nuclear Medicine and Molecular Imaging | 19 | 7 |
| Caroli A | 2007 | 15010013 | Journal of Neurology | 14 | 2 |
| Bai F | 2008 | 20100037 | Neuroscience Letters | 20 | 7 |
| Long Z | 2016 | 20110048 | Neuroscience | 29 | 3 |
| Morbelli S | 2010 | 20120049 | European Journal of Nuclear Medicine and Molecular Imaging | 9 | 3 |
| Yin C | 2014 | 21010001 | Translational Neuroscience | 11 | 5 |
| Zhao Z | 2014 | 21010002 | BioMed Research International | 18 | 7 |

*For each experiment included from the BrainMap VBM database, the identification number (BrainMap ID) was provided.* ***Subj*** *= number of subjects in the clinical group;* ***Foci*** *= number of x-y-z coordinates of gray matter alteration (i.e., Controls > Patients).*

*Table S2.* Experiments included in the meta-analysis: Unrelated cognitive domains dataset.

| ***First author*** | ***Year*** | ***BrainMap ID*** | ***Journal*** | ***Subj*** | ***Foci*** |
| --- | --- | --- | --- | --- | --- |
| Martin R E | 2004 | 8020052 | Journal of Neurophysiology | 14 | 23 |
| Leveroni C L | 2000 | 30043 | Journal of Neuroscience | 11 | 2 |
| Menon V | 2001 | 30356 | Human Brain Mapping | 14 | 4 |
| Hagen M C | 2002 | 4040027 | Journal of Neurophysiology | 12 | 4 |
| Valentin V V | 2009 | 10050113 | Journal of Neurophysiology | 17 | 2 |
| Mathiak K | 2006 | 9090135 | Human Brain Mapping | 13 | 8 |
| Nelson A J | 2004 | 4040038 | Cognitive Brain Research | 6 | 1 |
| Kadosh R C | 2005 | 14050077 | Neuropsychologia | 15 | 5 |
| Farb N A S | 2012 | 16030026 | Social Cognitive and Affective Neuroscience | 16 | 7 |
| Pelletier M | 2003 | 6040032 | Neuroreport | 9 | 8 |
| Li G | 2003 | 30294 | Neuroreport | 20 | 4 |
| Schaefer M | 2007 | 12100078 | Brain Research | 14 | 1 |
| Petrini K | 2011 | 18080092 | PLoS ONE | 16 | 12 |
| Bueti D | 2008 | 8080196 | Journal of Cognitive Neuroscience | 14 | 5 |
| Dichter G S | 2007 | 7040095 | NeuroImage | 17 | 2 |
| Servos P | 2002 | 7070189 | Cerebral Cortex | 12 | 2 |
| Dionne J K | 2010 | 16030074 | Human Brain Mapping | 10 | 1 |
| Rothemund Y | 2007 | 7110310 | NeuroImage | 13 | 1 |
| Ramirez-Ruiz B | 2008 | 15010005 | Movement Disorders | 10 | 1 |
| Kuhtz-Buschbeck J P | 2005 | 7080218 | Journal of Urology | 22 | 6 |

*For each experiment included from the BrainMap functional database, the identification number (BrainMap ID) was provided.* ***Subj*** *= number of subjects in the experimental group;* ***Foci*** *= number of x-y-z coordinates of gray matter activation.*

***Figure S1.*** Comparison between the ALE map with canonical FWE thresholding and the BF map thresholed at log10(mBF)=5. Red voxels were present in both maps, blue voxels were present in the mBF map only, yellow voxels were present in the FWE map only. Values in bracket refers to the number of experiments in the dataset. Axial slices were obtained at the z coordinate reported.

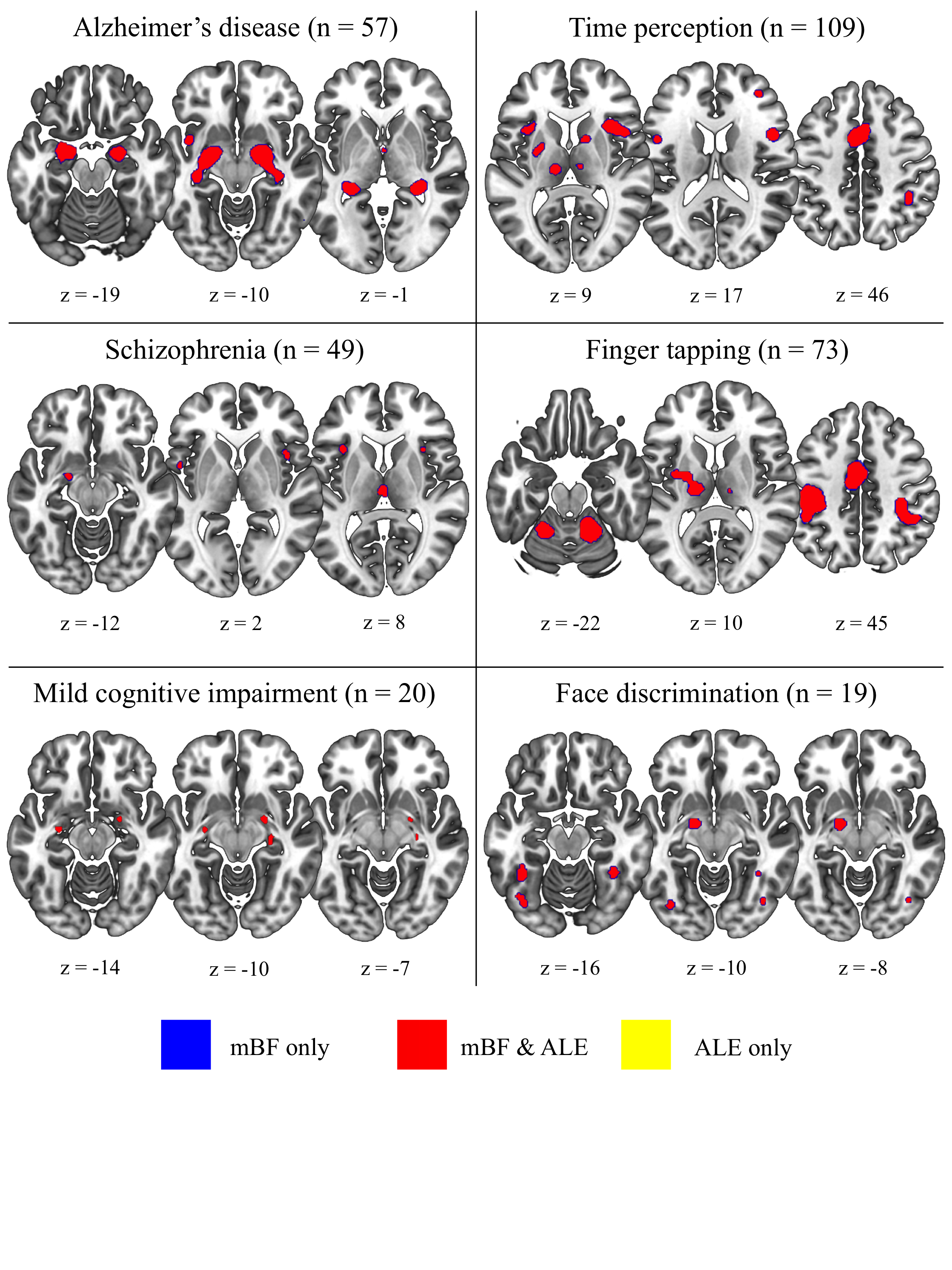

***Figure S2.*** Comparison between the ALE map with canonical c-FWE thresholding and the BF map thresholed at log10(mBF)=2. Red voxels were present in both maps, blue voxels were present in the mBF map only, yellow voxels were present in the FWE map only. Values in bracket refers to the number of experiments in the dataset. Axial slices were obtained at the z coordinate reported.

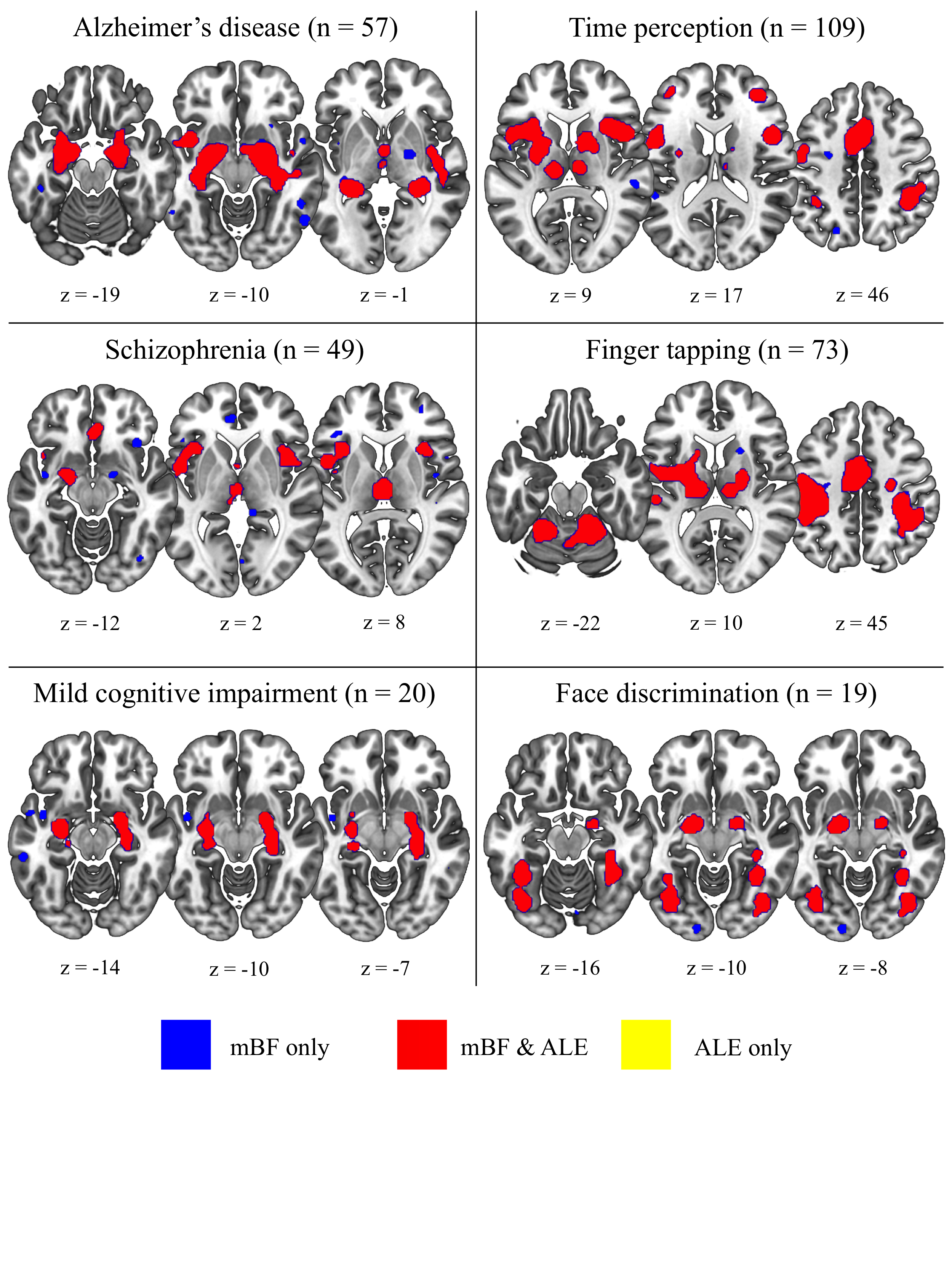

***Figure S3.*** Number of non-zero voxels appearing in the mBF maps obtained for the unrelated cognitive domains dataset (blue line) and the simulated datasets (red and yellow lines). The x axis was limited at log10(mBF)=4 since that threshold produced empty maps in each of the three conditions.

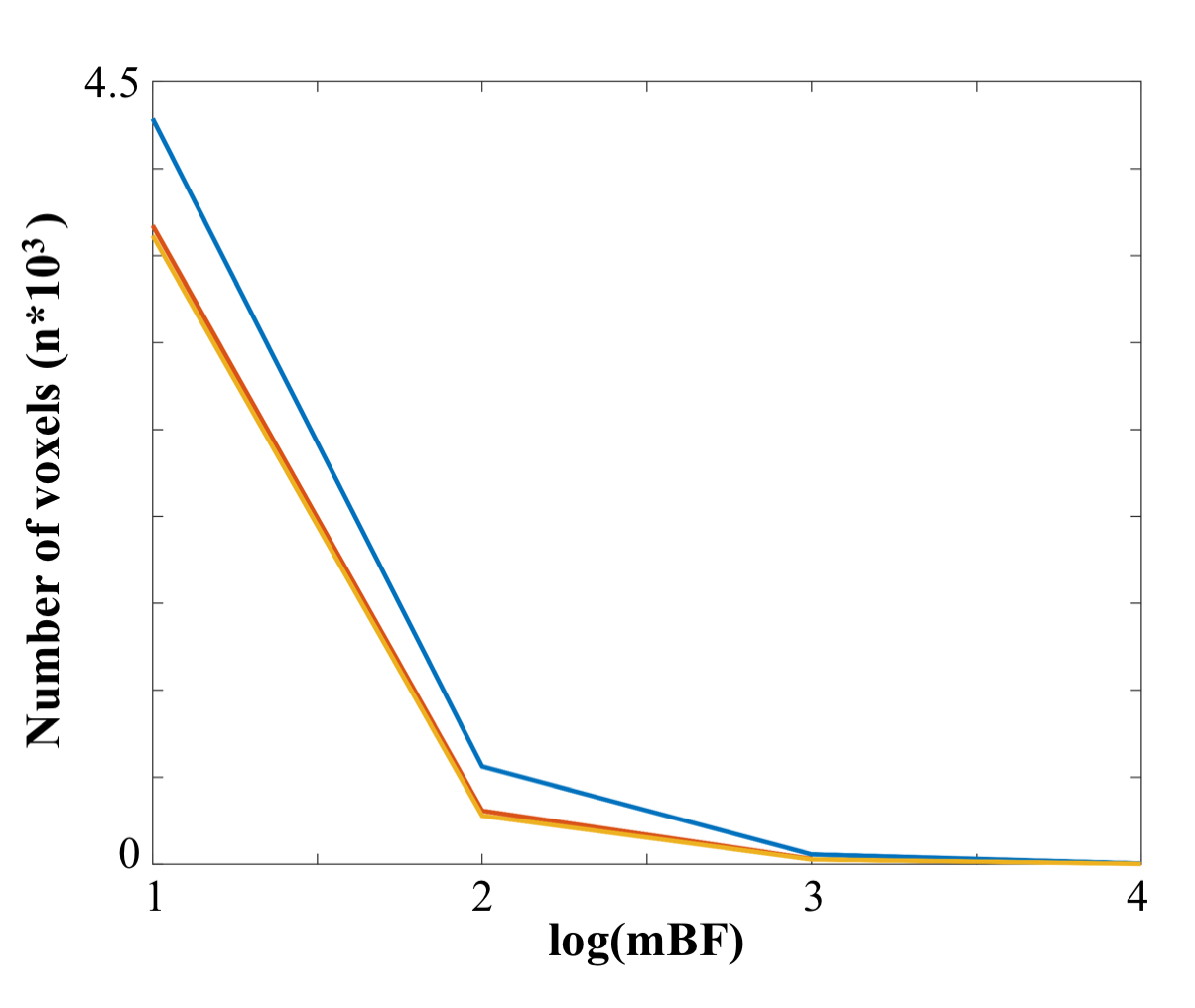
